# Checks and balances of the RTG pathway under arginine deprivation and canavanine exposure in *Saccharomyces cerevisiae*

**DOI:** 10.1101/2022.02.21.481281

**Authors:** Marina E. Druseikis, Shay Covo

**Affiliations:** Department of Plant Pathology and Microbiology, Robert H. Smith Faculty of Agriculture, Food and Environment. The Hebrew University of Jerusalem. Rehovot 76100001, Israel

## Abstract

Decades of research in *Saccharomyces cerevisiae* underlie the current dogma of mitochondrial retrograde (RTG) signaling: Rtg2-dependent translocation of the heterodimer Rtg1/Rtg3 from the cytoplasm to the nucleus induces the transcription of RTG-target genes under glutamate starvation or loss of respiration. We previously found that RTG mutants show severe growth inhibition from arginine deprivation and are highly sensitive to canavanine when grown on glucose. Here, we show that on solid media, RTG mutants are also sensitive to thialysine, a toxic lysine analog, although lysine deprivation causes a milder growth defect. Growth on an alternative carbon source restores RTG mutants’ ability to grow without arginine or lysine and improves their tolerance of toxic analogs; deletion of *MIG1* affords a similar rescue on glucose and improves canavanine tolerance, except for in *rtg2Δ*. It is well known that the target of rapamycin (TOR) signaling pathway inhibits the RTG pathway. Batch growth experiments with or without TOR inhibition reveal phenotypic and regulatory differences between RTG mutants. *rtg1Δ* can sustain simultaneous canavanine exposure and TOR inhibition via rapamycin, but *rtg2Δ* and *rtg3Δ* cannot. Surprisingly, our data show that under fermentative lifestyle and arginine deprivation, both RTG signaling and TOR activity are required. This expands the universe of TOR and RTG signaling, suggesting bilateral communication rather than unidirectional RTG regulation by TOR. To the best of our knowledge, this work shows for the first time that Rtg3 activity can be separate from its role as a heterodimer with Rtg1. This work also strongly suggests a specific role for Rtg2 in canavanine tolerance.

## Introduction

RTG genes (*RTG1, RTG2, RTG3*, and *MKS1*) encode for regulators of the mitochondrial ReTroGrade (RTG) pathway, which is activated in response to various physiological stresses, including loss of respiration capacity and amino acid depletion (1, 2). Rtg2 is the central regulator of the RTG pathway - it senses mitochondrial dysfunction and transduces a signal through Rtg1/Rtg3. Rtg1 and Rtg3 work together as a heterodimer; when the retrograde response is activated, Rtg1/Rtg3 is translocated from the cytosol to the nucleus in an *RTG2*-dependent manner, where they induce transcription of RTG-target genes. In the absence of *RTG2*, Rtg1/Rtg3 remains in the cytoplasm. However, this *RTG2*-requirement for nuclear localization can be overridden by simultaneous deletion of *MKS1*. While Mks1 negatively regulates Rtg1/Rtg3 nuclear localization (3), binding of Rtg2 to Mks1 relieves this inhibition (4). Whereas *rtg1-3Δ* mutants are glutamate auxotrophs, both *mks1Δ* and *rtg2Δmks1Δ* strains exhibit phenotypes that are associated with RTG pathway activation: glutamate prototrophy and high expression of *CIT2*, the prototypical RTG-response gene (3, 5).

Although Rtg1 and Rtg3 act together as a single transcription factor, localization experiments with fluorescently-labeled Rtg1, Rtg2, and Rtg3 show that Rtg1 also has a function as a negative regulator of the RTG pathway, since it has a role in maintaining Rtg3 in the cytosol (Rtg3 is nuclear in *rtg1Δ*) (6). In fact, *rtg1Δ* takes precedence over *rtg2Δ* when it comes to the localization of Rtg3; Rtg3 is cytoplasmic in *rtg2Δ*, but Rtg3 is nuclear in *rtg1Δrtg2Δ*.

It is still not understood how the cytoplasmic protein Rtg2 directs Rtg1/Rtg3 to the nucleus. Nonetheless, the Target of Rapamycin (TOR) is a known regulator of the RTG pathway (3). In yeast, TOR is composed of two similar but different proteins - TORC1, which responds to rapamycin, and TORC2, which is not rapamycin sensitive (7). TOR signaling is activated when conditions are pro-growth - namely sufficient nitrogen, glucose, and amino acids - to ensure there are enough supplies to carry out protein assembly (8, 9). Both transcription and translation are upregulated in response to TOR activation. TOR is a negative regulator of the RTG pathway through Mks1. When TOR is fully active, Mks1 is not bound to Rtg2 (instead it binds to Bmh1/2), and Rtg1/Rtg3 is cytoplasmic, meaning the RTG pathway is off (10). Upon TOR inhibition, Rtg2 binds Mks1, and the expression of RTG-target genes is increased, thus the RTG pathway is on (3).

Genes induced by RTG activation function upstream of alpha-ketoglutarate (αKG) production, including *CIT1* and *CIT2* (mitochondrial and peroxisomal citrate synthase, respectively), *DLD3* (2-hydroxyglutarate transhydrogenase), *ACO1* (mitochondrial aconitase), *IDH1* and *IDH2* (mitochondrial isocitrate dehydrogenase) (1). In respiratory-competent strains, some of these genes (*CIT1, ACO1, IDH1, IDH2*) are also under HAP (Heme Activator Protein) control. The HAP complex is a heme-activated, glucose-repressed transcriptional activator of respiratory gene expression (11–13). We previously found that strains missing a single RTG gene are highly sensitive to canavanine when grown on glucose lacking arginine (glucose (-)arginine), and that canavanine alters expression of RTG-target genes in both WT and petite cells (14). This suggests that canavanine either interferes directly with RTG signaling or upstream of the Rtg2/Mks1 association, and RTG mutant sensitivity is likely due to insufficient levels of metabolites required for arginine biosynthesis that are under RTG regulation.

Like many metabolic pathways, RTG activity is regulated by the carbon source (15, 16). Under partially respiratory conditions, like growth on galactose, both RTG proteins and the HAP complex are required for wild-type respiration. Under fully respiratory conditions, the HAP complex is still needed, but RTG proteins are not essential for respiration (11). In yeast, many metabolic and respiration-related genes are repressed when glucose is present. Glucose repression is mediated through the SNF1/AMPK pathway; when glucose is present, *MIG1* inhibits expression of some respiratory, gluconeogenic, and alternative carbon source genes (17). As glucose becomes limited, Snf1 inhibits Mig1, allowing expression of glucose-repressed genes; deletion of *MIG1* at least partially relieves glucose repression. It’s been shown that *CIT2* transcription requires the Snf1-dependent transcription factors *ADR1* and *CAT8* (16). Since Snf1 controls *ADR1* and *CAT8* through regulation of Mig1 and the corepressor Tup1-Cyc8 complex (18), it’s possible that glucose influences RTG-target gene transcription through Snf1 (19).

The TOR and SNF1/AMPK pathways are often considered opposing regulators of downstream processes like amino acid biosynthesis, nitrogen catabolite repression, and autophagy (20, 21). Under nutrient-rich conditions, TOR promotes growth and suppresses autophagy; under starvation conditions (when SNF1/AMPK is active), TOR is inhibited and autophagy is induced, which provides substrates for rebuilding and repairing cells. The seemingly opposite roles of TOR and SNF1/AMPK come from studies in which cells are either in nutrient-rich or nutrient-starved conditions (22); the nuances of these two pathways in maintaining homeostasis during metabolic fluctuations are not well understood.

Here we exposed RTG mutants and mutants of their downstream targets to low doses of canavanine and thialysine, toxic analogs of arginine and lysine, respectively. Under arginine or lysine deprivation conditions, even low-dose analog exposure causes a strong demand for the respective amino acid biosynthesis and thus allow us to investigate the role of each of the RTG components in high resolution. We found that Mig1 inhibits RTG-alternative mechanisms of arginine and lysine biosynthesis, and that Rtg1, Rtg2, and Rtg3 differ in the way they are regulated by Mig1, TOR signaling, and canavanine. These results expand the universe of the RTG pathway in agreement with recent findings (23, 24).

## Materials and Methods

### Yeast Strains

We use the WT laboratory strain BY4741; *pif1Δ, mig1Δ, rtg1Δ, rtg2Δ, rtg3Δ, mks1Δ, cit1Δ, cit2Δ, dld3Δ, idh1Δ*, and *idh2Δ* are also of the BY4741 background and from the library constructed by Giaever et al. (25). Double knockouts were constructed by transforming one of the above G418-resistant mutants with a PCR product containing a *URA3* gene to replace the gene of interest (a RTG gene or RTG-target gene) and selection done on -URA, G418, and YPG media (to screen for respiratory status) followed by PCR confirmation. Primers used for transformation and confirmation can be found in Supplementary Table 1. A minimum of two independent transformants with matching phenotypes in growth experiments were verified for each RTG double mutant and RTG-target gene double mutant to reduce the likelihood that an observed phenotype is due to a random additional mutation. For *MIG1/RTG* double mutants, a minimum of three isolates were obtained and strains were confirmed to not be *CAN1* mutants.

### Media

All experiments with canavanine were carried out in complete synthetic media (CSM) without arginine ((-)arginine). Similarly, experiments using thialysine were carried out on media lacking lysine ((-)lysine). YPD solid media contains 2% Bacto Agar, 2% Bacto Peptone, 1% Bacto Yeast Extract (Difco, Sparks, MD, USA), and 2% anhydrous dextrose (Avantor Performance Materials, Center Valley, PA, USA) dissolved in DDW. For YPD + G418 plates, G418 was added to 400 ml for a final concentration of 8 μg/ml. All other media (complete, YPG, media lacking uracil, or media lacking a specific amino acid with or without a toxic amino acid analog) contain 2% Bacto Agar, 0.67% Yeast Nitrogen Base (YNB) without Amino Acids (Difco Laboratories), 0.082% Complete Supplement Mixture (no amino acid drop-out) or 0.074% Complete Supplement Mixture Drop-out: ARG, LYS, or URA (Formedium). All media (excluding YPD) are brought to pH 5.8 using NaOH pearls (Bio-Lab LTD, Jerusalem, IL, USA), autoclaved, and 2% glucose, 2% galactose, or 3% lactic acid added after autoclaving. Liquid medias were prepared in the same fashion as solid media with the exclusion of Bacto Agar.

### Growth assays

For growth assays on solid media (spot assays), using a 96-well tissue culture plate (Jet Biofil), a single colony was placed in 100 μl DDW and underwent a tenfold serial dilution. Using a 6-by-8-column metal prong, approximately 1 μl from each well is pronged onto desired media. After allowing suspended cells to fully absorb into the plate, the plate is put into 30°C and growth photographed at the indicated time points.

For the 72-hour batch growth assay in liquid media, a single colony was grown overnight in 2 ml of YPD starter culture. All four types of media were prepared from the same stock of glucose (-)arginine, which was aliquoted into 50 ml tubes. The appropriate amount of stock solution of canavanine or rapamycin was added to glucose (-)arginine for a final concentration of 1 μg/ml canavanine, 0.025 μg/ml rapamycin, or 1 μg/ml canavanine + 0.025 μg/ml rapamycin. From each of the four types of media, 1.95 ml was aliquoted to a 2 ml Eppendorf tube, for a total of four Eppendorfs for each isolate. OD_600_ of tenfold-diluted overnight cultures were read the following morning and each culture was diluted to an OD of 0.0375 in each Eppendorf. A total of 16 isolates were tested during an experiment, with eight isolates per 96-well plate for each type of media, for a total of eight plates (two plates per condition). For each isolate under each condition, 150 μl were taken from the corresponding 2 ml Eppendorf and moved to a predetermined well on the 96-well plate, for a total of 11 technical repeats per isolate per condition. Isolates are arranged in an alternating manner to control for geographical effects on growth. The first column of a plate always contained 150 μl of the corresponding “blank” media (media without cell culture) to ensure no contamination occurred. The eight 96-well culture plates were loaded into a 30°C incubator connected to the Tecan Spark^®^ plate reader. For 72 hours, every hour plates were robotically removed from the incubator and loaded onto the Tecan Spark^®^ where OD was read after agitation, and the plate returned to the incubator until the next measurement. Data was initially collected and run through a script using MATLAB to ensure no geographical effects occurred, and average growth curves for each isolate plotted including standard deviations. Data was exported to excel for construction of figure graphs. Raw data (OD for each time point and standard deviations) can be found in Supplementary Table 2.

## Results

### Canavanine sensitivity of RTG single and double mutants reveals the non-redundancy of each RTG gene and is influenced by the carbon source

We previously found that RTG mutants are highly sensitive to canavanine when grown on glucose (14), a carbon source that is preferentially fermented. When glucose is the sole carbon source, even a single mutation in the RTG pathway results in poor growth when arginine is lacking. To detangle the role of RTG genes in canavanine tolerance, we created a set of double RTG knockout mutants, each containing two deletions made from combinations of four genes of the RTG pathway: *rtg1Δ, rtg2Δ, rtg3Δ*, and *mks1Δ*. We then tested growth on glucose (-)arginine with a range of canavanine concentrations and found that all single and double RTG mutants show severe growth inhibition at just 0.25 μg/ml canavanine (a fraction of the dose that is toxic to another highly sensitive background - petites) (Fig. 1A, left panel), supporting the notion that all RTG genes work together. This extreme sensitivity also occurs in *mks1Δ* and *rtg2Δmks1Δ;* in both of these backgrounds, the RTG pathway is considered to be constitutively on, measured by high *CIT2* expression and the ability to grow without glutamate (5). In agreement with the literature, our *rtg2Δmks1Δ* strain does show better growth than other RTG mutants when grown on media lacking arginine, but *mks1Δ* does not; however, when canavanine is added, *mks1Δ* and *rtg2Δmks1Δ* are just as sensitive as the other *rtgΔ* mutants. This suggests that although the RTG pathway is turned on, it is not functional at either WT-level or petite-level in these two backgrounds under these conditions. Since RTG activity can be influenced by the carbon source, we tested canavanine sensitivity in two other carbon sources: galactose, which is used for both fermentation and respiration, and lactic acid, which is non-fermentable and essentially forces the cell to respire. Single and double RTG mutants show arginine auxotrophy when grown on glucose, but not on galactose or lactic acid (Fig. 1A). When grown on the non-fermentable carbon source lactic acid, RTG mutants can tolerate 0.25 – 0.5 μg/ml canavanine (Fig. 1A, Supp. Fig. 1), but at higher doses they are subject to the lactic acid effect, a phenomenon we previously reported in both WT and *can1Δ* mutants (14).

**Figure 1.**
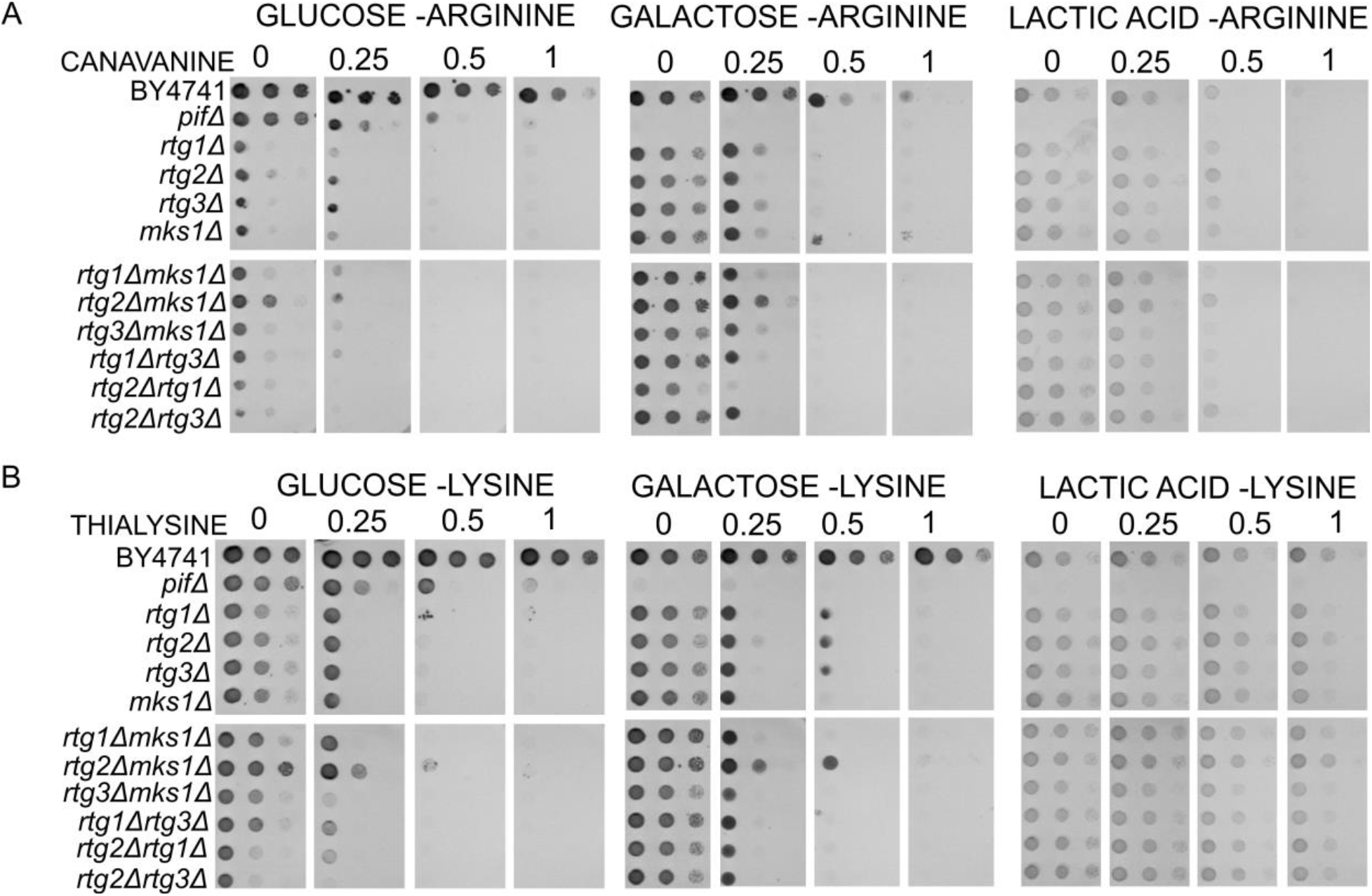
Alternative carbon sources improve growth of RTG mutants on (-)arginine and (-)lysine. WT BY4741, *pif1Δ*, and *rtgΔ* single and double mutants grown on A) glucose (-)arginine, galactose (-)arginine, and lactic acid (-)arginine with the indicated amount of canavanine (μg/ml); and B) glucose (-)lysine, galactose (-)lysine, and lactic acid (-)lysine with the indicated amount of thialysine (μg/ml). A single colony was put in 100 ul of DDW, serially diluted, pronged onto the indicated media using a 6X8 metal pronger, and grown in 30°C. Growth after 3 days.

Growth on galactose allows us to see differences in canavanine tolerance between the single and double RTG mutants: *rtg2Δrtg1Δ* is the only mutant that does not show growth on 0.25 μg/ml canavanine after three days; *rtg2Δ* shows a 10-fold reduction in growth in comparison with other RTG mutants, a difference that is not detected on glucose or lactic acid; and *rtg2Δmks1Δ* shows the best growth of all the mutants (Fig. 1A). This indicates that the role of each individual RTG gene in canavanine tolerance is not redundant and responds to the fermentation/respiration ratio. Given that RTG mutants can tolerate canavanine as well as WT when grown on lactic acid, our data show that the RTG pathway is not critical under strictly respiratory conditions.

### Thialysine tolerance is respiration-dependent in RTG mutants

To determine if the effect of the fermentation/respiration ratio of RTG mutants is specific to canavanine or more general, we grew RTG mutants on glucose (-)lysine media and exposed them to thialysine (AEC), a toxic amino acid analog of lysine. Notably, RTG mutants do not show poor growth on glucose media lacking lysine, although colony forming ability is decreased compared to growth on galactose (-)lysine or lactic acid (-)lysine (Fig. 1B). When grown on glucose or galactose for three days, all RTG mutants, except for *rtg2Δmks1Δ*, show severe growth inhibition at 0.25 μg/ml AEC and are unable to grow at 1 μg/ml AEC on either carbon source (Fig. 1B). After one week’s growth, RTG mutants show improved growth on galactose + AEC compared to glucose + AEC, like what we see in canavanine (Supp. Fig. 1). However, after only three days, all RTG mutants show sufficient growth on 1 μg/ml AEC when lactic acid is the carbon source (Fig. 1B). Unlike growth in canavanine, the AEC sensitivity assay does not detect sensitivity differences between individual *RTG* genes (Fig. 1, Supp. Fig. 1). Notably, we do not see a nuanced dependency on the fermentation/respiration ratio - strict respiration confers thialysine tolerance in RTG mutants.

### On solid media, relief of glucose repression via mig1Δ rescues rtg1Δ and mks1Δ from glucose-induced canavanine/thialysine sensitivity, but not rtg2Δ or rtg3Δ

We see that alternative carbon sources that require respiration improve, to a point, RTG mutants’ tolerance to both canavanine and thialysine; under such conditions, Mig1 is released from glucose repression and the cell can undergo transcription of respiratory-related genes (26). Thus, we wondered if relief of glucose repression via *MIG1* deletion could rescue RTG mutants from glucose-induced canavanine/thialysine toxicity. Interestingly, we observe different phenotypes between the four *mig1ΔrtgΔ* strains (Fig. 2). When grown on glucose (-)arginine/(-)lysine solid media, both *mig1Δrtg1Δ* and *mig1Δmks1Δ* can tolerate 1 μg/ml canavanine and 1 μg/ml AEC. This means *mig1Δ* rescues *rtg1Δ* in an RTG-independent way. Because *MKS1* is a negative regulator of the RTG pathway, without doing the triple KO of *mig1Δrtg1Δmks1Δ*, we are unable to say if *mig1Δ* rescues *mks1Δ* in an RTG-independent or RTG-dependent manner. More importantly, *mig1Δ* does not rescue *rtg2Δ* or *rtg3Δ* from canavanine/thialysine toxicity, although it does improve the ability of these strains to grow on media lacking arginine or lysine (Fig. 2); thus, on solid media, *RTG2* and *RTG3* are required for tolerance of glucose-induced canavanine/thialysine toxicity in a *MIG1*-independent manner.

**Figure 2.**
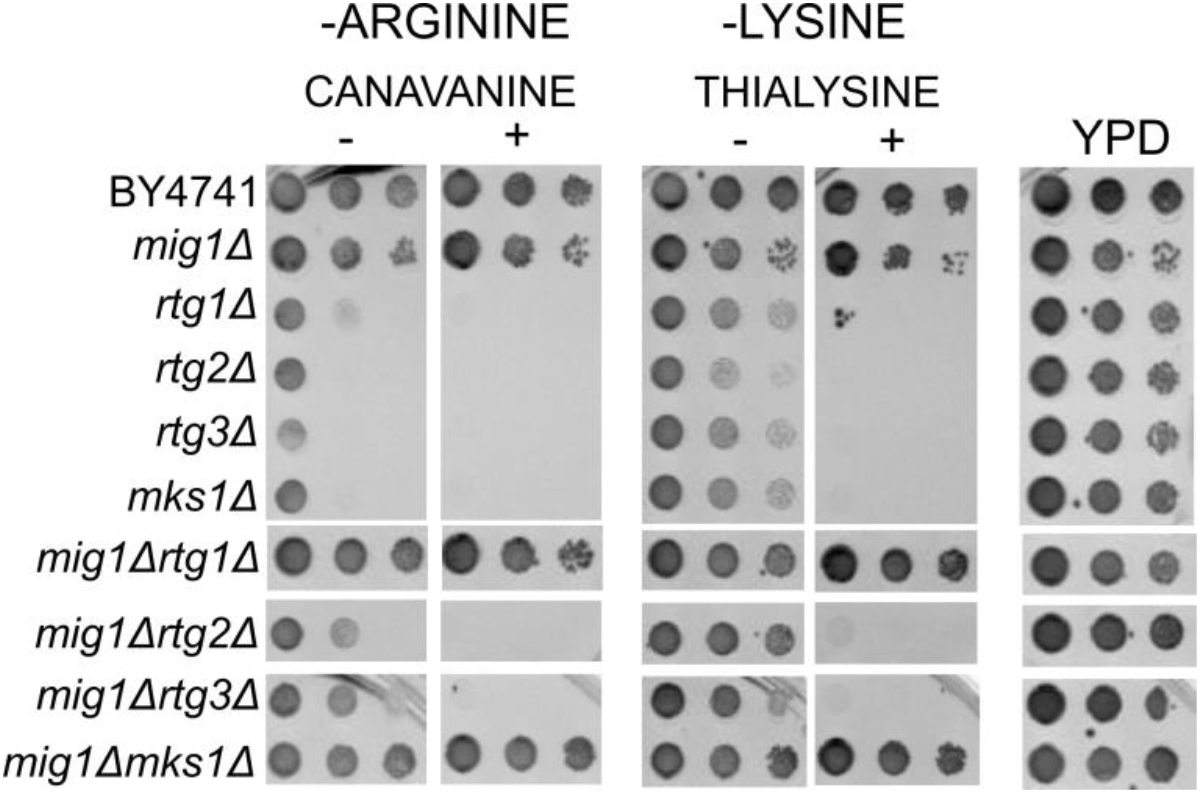
*MIG1* deletion improves growth of RTG mutants on (-)arginine and (-)lysine. WT BY4741, *rtgΔ* single mutants, and *mig1ΔrtgΔ* double mutants grown on glucose (-)arginine +/- canavanine 1 μg/ml, and glucose (-)lysine +/- thialysine 1 μg/ml, and YPD. A single colony was put in 100 ul of DDW, serially diluted, pronged onto the indicated media using a 6X8 metal pronger, and grown in 30°C. Growth after 4 days (glucose (-)arginine, glucose (-)lysine) and 2 days (YPD).

### Canavanine exposure in liquid media improves resolution of RTG1, RTG2, and RTG3 separation of function under arginine deprivation conditions in glucose

We grew strains in a 96-well plate in glucose (-)arginine with and without 1 μg/ml canavanine and measured growth (OD) for 72 hours. We note that, as we previously published, liquid media allows sufficient diffusion of nutrients that become limited in solid media. For this reason, the experiments done in liquid media show milder growth defects than those performed above. Growth in liquid glucose (-)arginine allows us to see in higher resolution what we see on plates; WT grows well, is the quickest to enter log phase, and has the highest OD at stationary phase (Fig. 3A). Compared to WT, the petite strain *pif1Δ* shows a slight increase in lag phase and reduced OD at stationary phase but maintains a similar log-phase growth rate. All RTG mutants show a reduced growth rate and further reduced OD (compared to *pif1Δ*) at stationary phase when glucose is present and arginine is lacking; we also detect a clear growth difference in *rtg2Δ* that is not present in the other mutants (Fig. 3A, 3B). When canavanine is added, there’s an elongation of lag phase in the WT, a characteristic of metabolic remodeling, and the final OD at stationary phase is the same as without drug. *pif1Δ* also undergoes an elongation of lag phase, although final OD is significantly reduced (Fig. 3A). Canavanine has a profound effect on the growth curves of RTG mutants; the log-phase growth rate is strongly reduced, and it’s unclear if cells have reached stationary phase at 72 hours, with the exception of *rtg2Δmks1Δ* (Fig. 3C; Supp. Fig. 2, Supp). However, even *rtg2Δmks1Δ* takes longer than *pif1Δ* to leave the lag phase (Supp. Fig. 3). Liquid media reveals a separation of phenotypes of the *RTG* genes under canavanine exposure even in glucose-repressing conditions: *rtg2Δ* shows the greatest growth inhibition, reflective of what we see on solid galactose with canavanine (Fig. 1A). We also detect a difference between *rtg1Δrtg3Δ* and *rtg2Δ* (Fig. 3C). Although these two strains have similar growth curves in glucose (-)arginine (Fig. 3B), in the presence of canavanine, *rtg2Δ* grows slower and reaches a lower final OD than *rtg1Δrtg3Δ* (Fig. 3C, Supp. Fig. 3). This phenotypic difference between *rtg1Δrtg3Δ* and *rtg2Δ* demonstrates that *RTG1, RTG2*, and *RTG3* have some non-redundant functions; turning off the RTG pathway via loss of the transcriptional regulator (Rtg1/Rtg3 heterodimer) has a lesser effect on growth than removing the central positive regulator, Rtg2. This again suggests that Rtg2 holds some additional function under these conditions, perhaps as a chaperone to other proteins required for canavanine tolerance, given Rtg2 is largely cytoplasmic. Nevertheless, when it comes to canavanine tolerance, this alternative role of Rtg2 is still secondary to its classic role in RTG pathway activation, since *rtg2Δmks1Δ*, which has the pathway constitutively on, grows better than *rtg1Δrtg3Δ* (Fig. 3C, Supp. Fig. 3).

**Figure 3.**
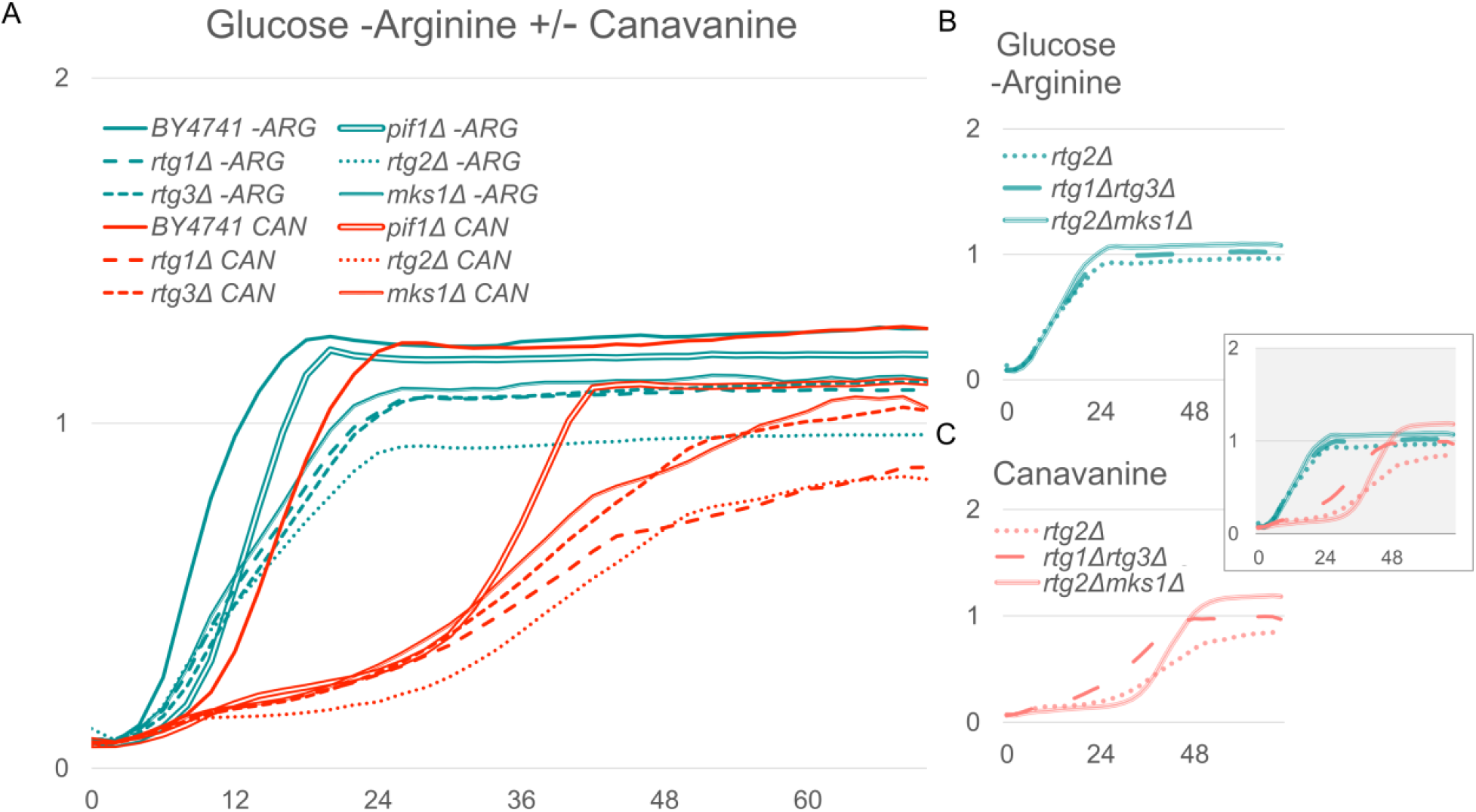
RTG-dependent growth in glucose (-)arginine and the effect of canavanine. A) Growth curves of WT BY4741, *pif1Δ*, and *rtgΔ* single mutants grown in liquid glucose (-)arginine (teal) and glucose (-)arginine + 1 μg/ml canavanine (red); B) Growth curves of *rtg2Δ, rtg1Δrtg3Δ*, and *rtg2Δmks1Δ* in glucose (-)arginine; C) Growth curves of *rtg2Δ, rtg1Δrtg3Δ*, and *rtg2Δmks1Δ* in glucose (-)arginine + 1 μg/ml canavanine. Graph imposed on B) and C) shows B) and C) on the same graph. X-axis = hours, Y-axis = OD_600_.

In comparison, relief of glucose repression via deletion of *MIG1* has no negative effect on WT growth in glucose (-)arginine with or without canavanine. In the RTG mutant backgrounds grown in liquid glucose (-)arginine, deletion of *MIG1* in *rtg1Δ* and *mks1Δ* restores growth to WT levels, has no effect on *rtg3Δ*, and only slightly increases final OD at stationary phase in *rtg2Δ* (Fig. 4A). In the presence of canavanine, deletion of *MIG1* restores *rtg1Δ, rtg3Δ*, and *mks1Δ* growth curves to the usual “S” shape - that is, lag, log, and stationary phase are all clearly discernible (Fig. 4B). In other words, canavanine has little effect on the growth rate of *mig1Δrtg1Δ, mig1Δmks1Δ*, and *mig1Δrtg3Δ* (Supp. Fig. 2); however, canavanine has a profound effect on the growth curve of *mig1Δrtg2Δ* (Fig. 4B, Supp. Fig. 2). Thus, deletion of *MIG1* in the *rtg2Δ* background doesn’t improve growth in canavanine. While the observed tolerance from relief of glucose repression is RTG-independent (based on *mig1Δrtg1Δ* and *mig1Δrtg3Δ* showing improved growth in liquid culture + canavanine compared to *rtg1Δ* and *rtg3Δ*), it appears to be RTG2-dependent, since relief of glucose repression via *MIG1* deletion has no benefit in the absence of Rtg2 (Figs. 2, 4B; Supp. Fig. 2).

**Figure 4.**
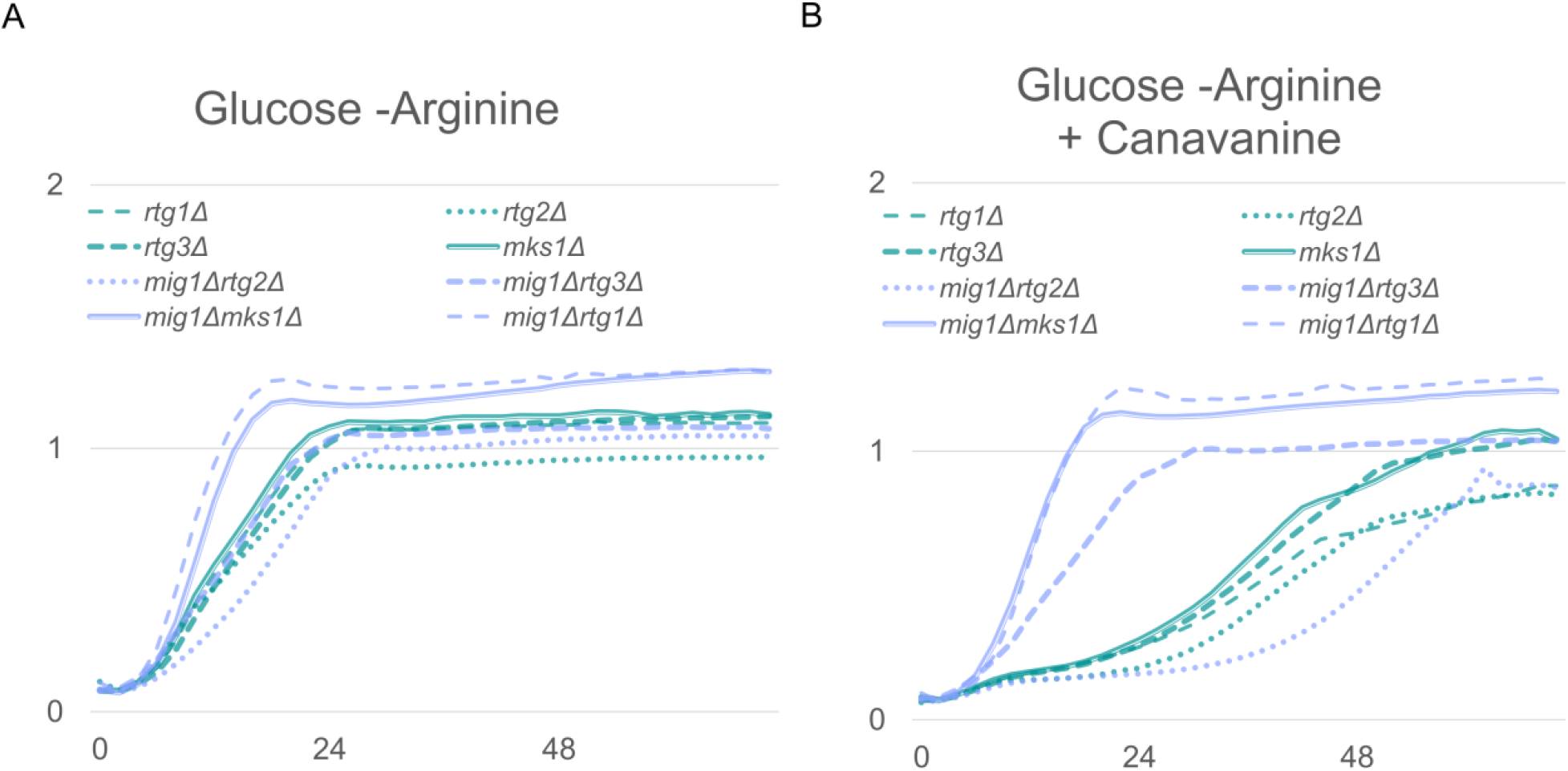
*MIG1* deletion restores growth in RTG mutants exposed to canavanine except for *rtg2Δ*. A) Growth curves of *rtgΔ* single mutants (teal) and *mig1ΔrtgΔ* double mutants (purple) grown in liquid glucose (-)arginine; B) Growth curves of *rtgΔ* single mutants (teal) and *mig1ΔrtgΔ* double mutants (purple) grown in liquid glucose (-)arginine + 1 μg/ml canavanine. X-axis = hours, Y-axis = OD_600_.

### Under glucose-repressing conditions, a fully functional RTG pathway is needed when TORC1 is inhibited

It is well documented that rapamycin inhibition of TORC1 causes RTG2-dependent nuclear accumulation of Rtg1/Rtg3, which turns on the RTG pathway and induces expression of RTG*-*target genes. Thus, we exposed WT, *pif1Δ, mig1Δ*, RTG mutants, *rtg1Δrtg3Δ, rtg2Δmks1Δ*, and *mig1ΔrtgΔ* double mutants to a sublethal dose of rapamycin (0.025 μg/ml). Upon rapamycin exposure, WT, *pif1Δ*, and *mig1Δ* have growth curves that are strikingly similar to their respective growth curves on glucose (-)arginine (Supp. Fig. 2). For RTG mutants, relative to WT, log-phase growth rate is reduced by rapamycin exposure, as well as final OD at stationary phase (Fig. 5A, Supp. Fig. 2). This expands the current understanding of the relationship between TOR and the RTG pathway. It’s known that under nutrient-replete conditions, TOR negatively regulates RTG pathway activation (TOR on, RTG off), and rapamycin inhibition of TOR induces nuclear localization of Rtg1/Rtg3 and subsequent RTG-target gene expression (TOR off, RTG on). We demonstrate that under arginine deprivation in glucose, there is RTG-dependent growth, and there is TOR-dependent growth in an RTG-independent manner (Supp. Fig. 2); both the RTG pathway and TOR activity are required for WT-level growth in glucose lacking arginine. If RTG signaling is absent, inhibition of TORC1 results in a further reduction in growth (Figs. 5A, 6; Supp. Fig. 2).

**Figure 5.**
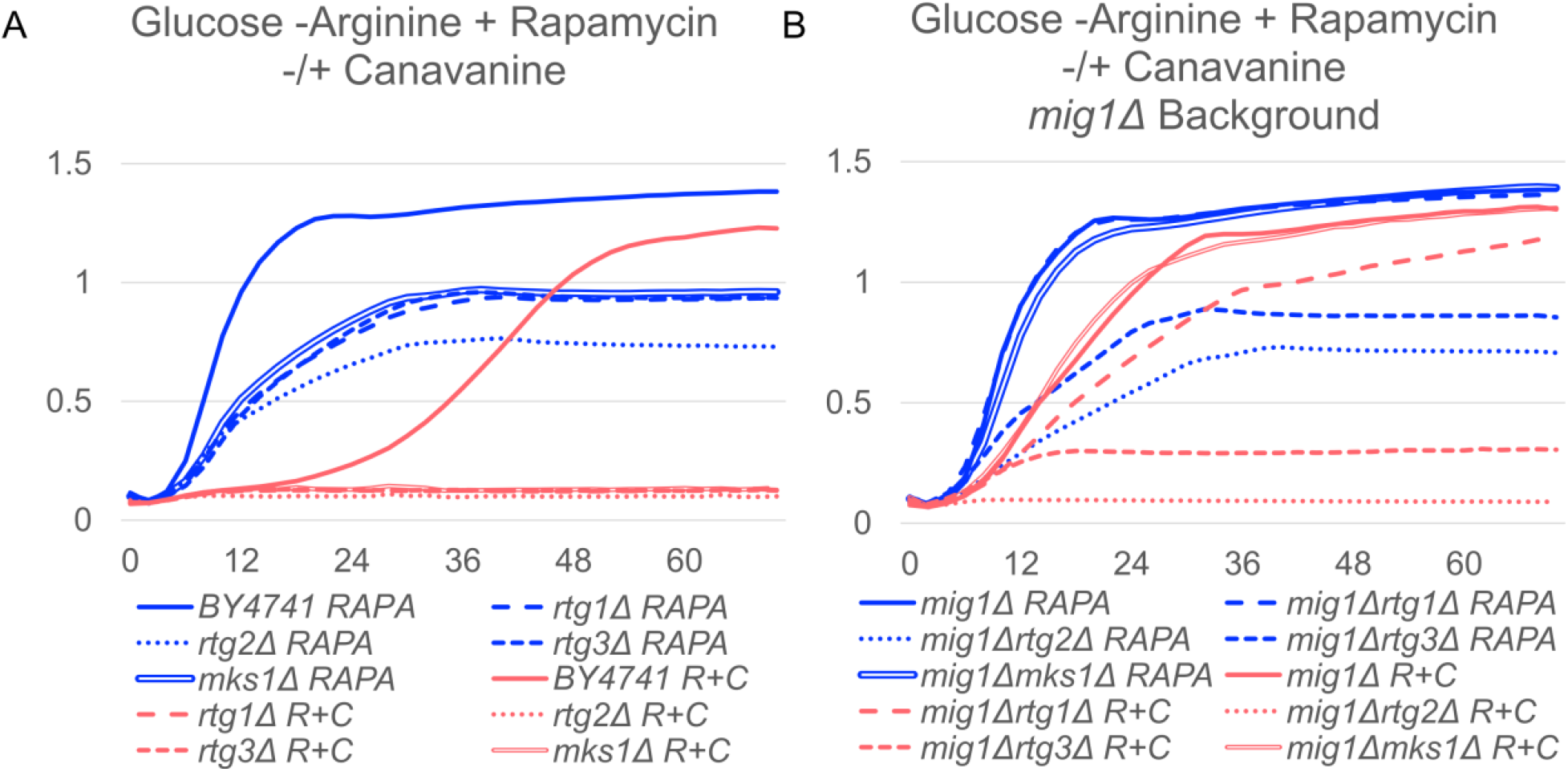
Both RTG activity and TOR activity are required for growth under arginine deprivation. A) Growth curves of BY4741 and *rtgΔ* single mutants in liquid glucose (-)arginine + 0.025 μg/ml rapamycin (blue) and glucose (-)arginine + 0.025 μg/ml rapamycin + 1 μg/ml canavanine (red). B) Growth curves of *mig1Δ* and *mig1ΔrtgΔ* single mutants in liquid glucose (-)arginine + 0.025 μg/ml rapamycin (blue) and glucose (-)arginine + 0.025 μg/ml rapamycin + 1 μg/ml canavanine (red). X-axis = hours, Y-axis = OD_600_.

**Figure 6.**
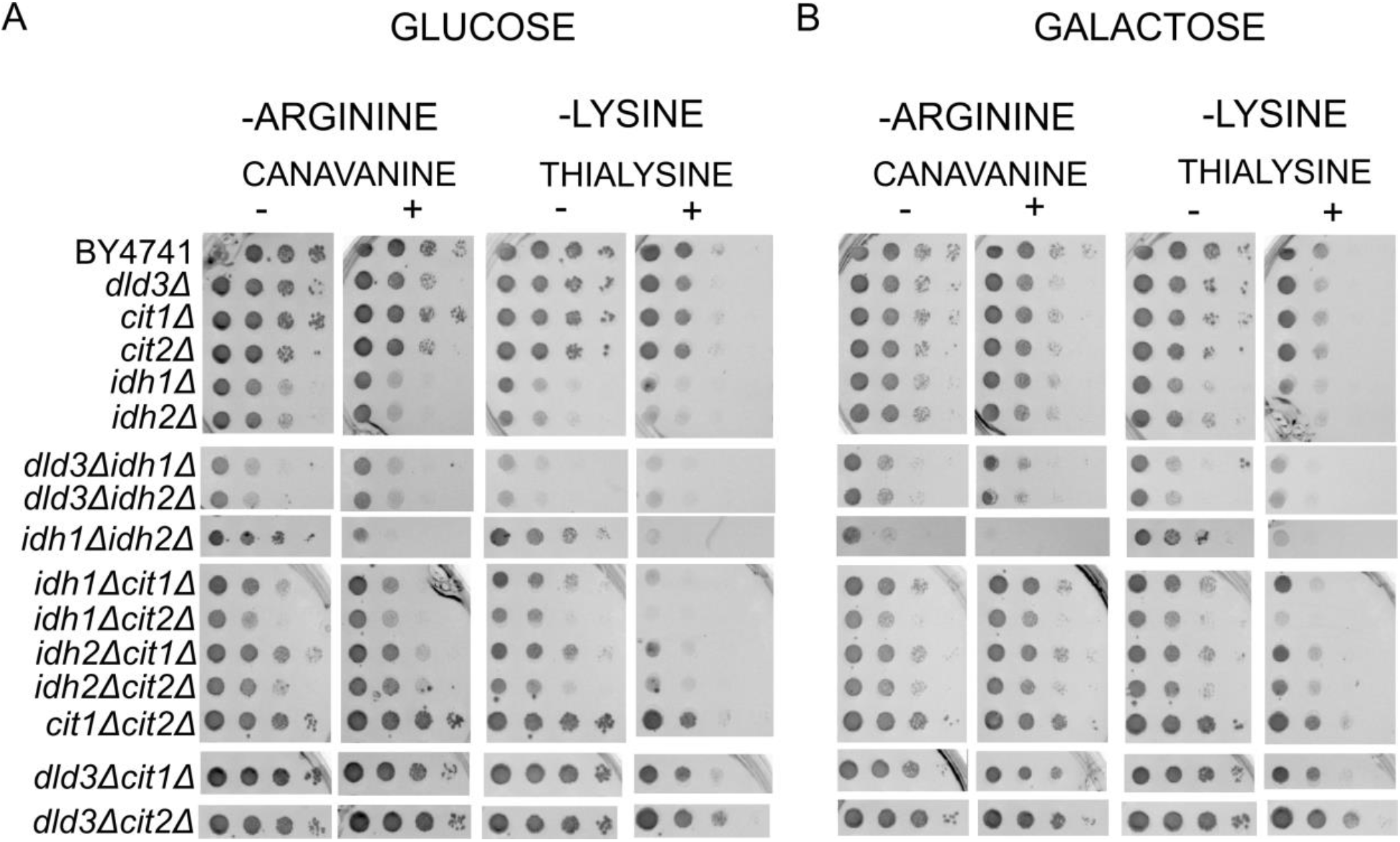
RTG-target genes directly involved in aKG levels show growth inhibition in (-)arginine and (-)lysine. Spot assay of WT BY4741 and RTG-target gene single and double mutants grown on A) Glucose (-)arginine +/- 1 μg/ml canavanine and glucose (-)lysine +/- 1 μg/ml thialysine; B) Galactose (-)arginine +/- 1 μg/ml canavanine and galactose (-)lysine +/- 1 μg/ml thialysine. A single colony was put in 100 μl of DDW, serially diluted, pronged onto the indicated media using a 6X8 metal pronger, and grown in 30°C. Growth after 3 days.

### TORC1 and RTG2/RTG3 are required for canavanine tolerance in glucose

TOR is activated when conditions are good - like when there are sufficient amounts of nitrogen, glucose, and amino acids - which greenlights anabolic processes like ribosome biogenesis, nucleotide synthesis, and translation. Thus, we wondered if TOR is needed for canavanine tolerance under our conditions: when there is sufficient nitrogen and glucose is abundant, but arginine is lacking. To answer this, we looked at the growth of our strains when simultaneously exposed to both rapamycin 0.025 μg/ml and canavanine 1 μg/ml. Our collection of RTG and *mig1Δ* mutants reveals nuances of the nutrient sensing crosstalk between TOR and the RTG pathway. Under the double exposure of rapamycin and canavanine, both WT and *pif1Δ* have an elongated lag phase, although *pif1Δ’s* lag phase is notably longer than WT’s (Supp. Fig. 2). *mig1Δ* shows the greatest tolerance; although the log-phase growth rate is reduced, there is no elongation of lag phase, and final OD is high (Supp. Fig. 2, Fig. 5B). The combined exposure of rapamycin and canavanine is lethal for all single and double RTG mutants tested - including both *mks1Δ* and *rtg2Δmks1Δ*, suggesting that the crosstalk between TOR and RTG may involve another layer of amino acid sensing/regulation since the RTG pathway should be constitutively on in these backgrounds, or that canavanine’s effect on the RTG pathway is modulated through TOR. However, relief of glucose repression via deletion of *MIG1* rescues *mks1Δ* as well as *rtg1Δ:* these double mutants grow as well as WT (defined by log-phase growth rate and final OD at stationary phase), but forgo the metabolic remodeling observed in WT and *pif1Δ* (Fig. 5B, Supp. Fig. 2). *rtg2Δmig1Δ* and *rtg3Δmig1Δ* do not show the same rescue (Fig. 5B, Supp. Fig. 2). Taken together, this means TORC1 is required for canavanine tolerance in an RTG-independent manner. This can be bypassed by relief of glucose repression (via *mig1Δ* deletion), but is still *RTG2/RTG3*-dependent; in other words, both TORC1 and *RTG2/RTG3* are required for canavanine tolerance when grown on glucose. Supporting this notion is a reported physical interaction between Rtg3 and Kog1 (27), a subunit of the TORC1 complex that has recently been shown to play a role in balancing carbon flux toward amino acid biosynthesis and gluconeogenesis (28).

### RTG-target genes directly involved in maintaining alpha-ketoglutarate levels show the most severe growth defects on (-)arginine and (-)lysine

To further investigate how the RTG pathway is needed in both canavanine and thialysine tolerance, we created another set of double mutants in which we combined pairs of gene deletions of several well-studied RTG-target genes: *CIT1, CIT2, IDH1, IDH2*, and *DLD3*. We were unable to isolate an *aco1Δ* mutant in the BY4741 background; multiple transformation attempts in our lab were unsuccessful, and BY4741 *aco1Δ* from the knockout collections of three other independent labs were confirmed via PCR to contain the *ACO1* gene, while simultaneously not showing the petite phenotype. In total, we created nine RTG-target gene double mutants. We previously reported that of the parent single mutants, only *idh1Δ* and *idh2Δ* show an arginine biosynthesis defect on glucose (-)arginine and glucose (-)lysine (29). Interestingly, *idh1Δ* and *idh2Δ* show a greater growth defect on glucose (-)lysine (Fig. 6), whereas RTG mutants show a greater growth defect on glucose (-)arginine (Fig. 1). In both *idh1Δ* and *idh2Δ*, growth defects on glucose (-)arginine and glucose (-)lysine are absent when grown on galactose (Fig. 6), similar to RTG mutants. The combined mutations of *dld3Δidh1Δ* and *dld3Δidh2Δ* show greater arginine/lysine biosynthesis deficits than the respective single mutants; these two mutants along with *idh1Δidh2Δ* also show synthetic lethality on both canavanine and thialysine (Fig. 6). The ability of a double knockout to tolerate canavanine does not mean it will be able to tolerate thialysine equally well - *idh1Δcit1Δ, idh1Δcit2Δ, idh2Δcit1Δ*, and *idh2Δcit2Δ* are more sensitive to thialysine than canavanine when grown on glucose (Fig. 6). In RTG mutants, growth on galactose has a clear positive effect on growth both with and without drug under arginine deprivation and lysine deprivation. In the RTG-target gene double mutants, carbon source has less of an effect on toxic amino acid analog tolerance, although RTG-target gene mutants are far less sensitive to toxic analogs than RTG mutants (on glucose, RTG-target genes mutants can tolerate 1 μg/ml canavanine/thialysine, RTG mutants can barely tolerate 0.25 μg/ml canavanine/thialysine).

Considering these differences in growth and sensitivity between RTG mutants and RTG-target gene mutants, it is possible that some of the phenotypes we see in RTG mutants are a result of reduced αKG. Nonetheless, this demonstrates that under arginine deprivation, the loss of a single RTG protein is more detrimental than the loss of the two RTG-target genes, especially when combined with canavanine exposure (Fig. 1A, Fig. 6). However, under lysine deprivation, loss of *idh1Δ, idh2Δ*, or the combination of *dld3Δ* with one of these deletions has a greater inhibitory effect on growth than loss of a single RTG protein, even with galactose as the carbon source (Fig. 1B, Fig. 6). This suggests that under arginine deprivation on glucose, RTG proteins are required for more than just activation of RTG-target genes; under lysine deprivation, the phenotype we see in RTG mutants is likely a result of reduced αKG/glutamate.

## Discussion

In this work we study the growth of RTG mutants, the effect of inhibiting their regulators from both the SNF1/AMPK pathway and TOR, and growth of mutants in their downstream targets (RTG-target genes) under conditions of either arginine deprivation or lysine deprivation. We also expose yeasts to sublethal doses of the toxic amino acid analogs canavanine and thialysine; our hypothesis is that the toxic analogs increase the demand for amino acid biosynthesis of the respective deprived amino acid (canavanine increases arginine biosynthesis demand, and thialysine increases lysine biosynthesis demand). This hypothesis is based on the current understanding of the RTG pathway in metabolism, as well as our previous findings regarding low-dose canavanine exposure in WT and petite mutants (14). Thus, the simplest way to explain the phenotypes of RTG mutants is their inability to produce sufficient glutamate to meet demands for arginine or lysine deprivation. However, when we consider all the data, several results suggest a more complicated system, addressed below. We also note that in this work, we did not address other potential consequences of low-dose analog exposure, such as protein misfolding. Importantly, our results show that under arginine deprivation, both the RTG pathway and TOR are required for WT-level growth (Fig. 7). This expands the current understanding of the relationship between RTG signaling and TOR signaling.

**Figure 7.**
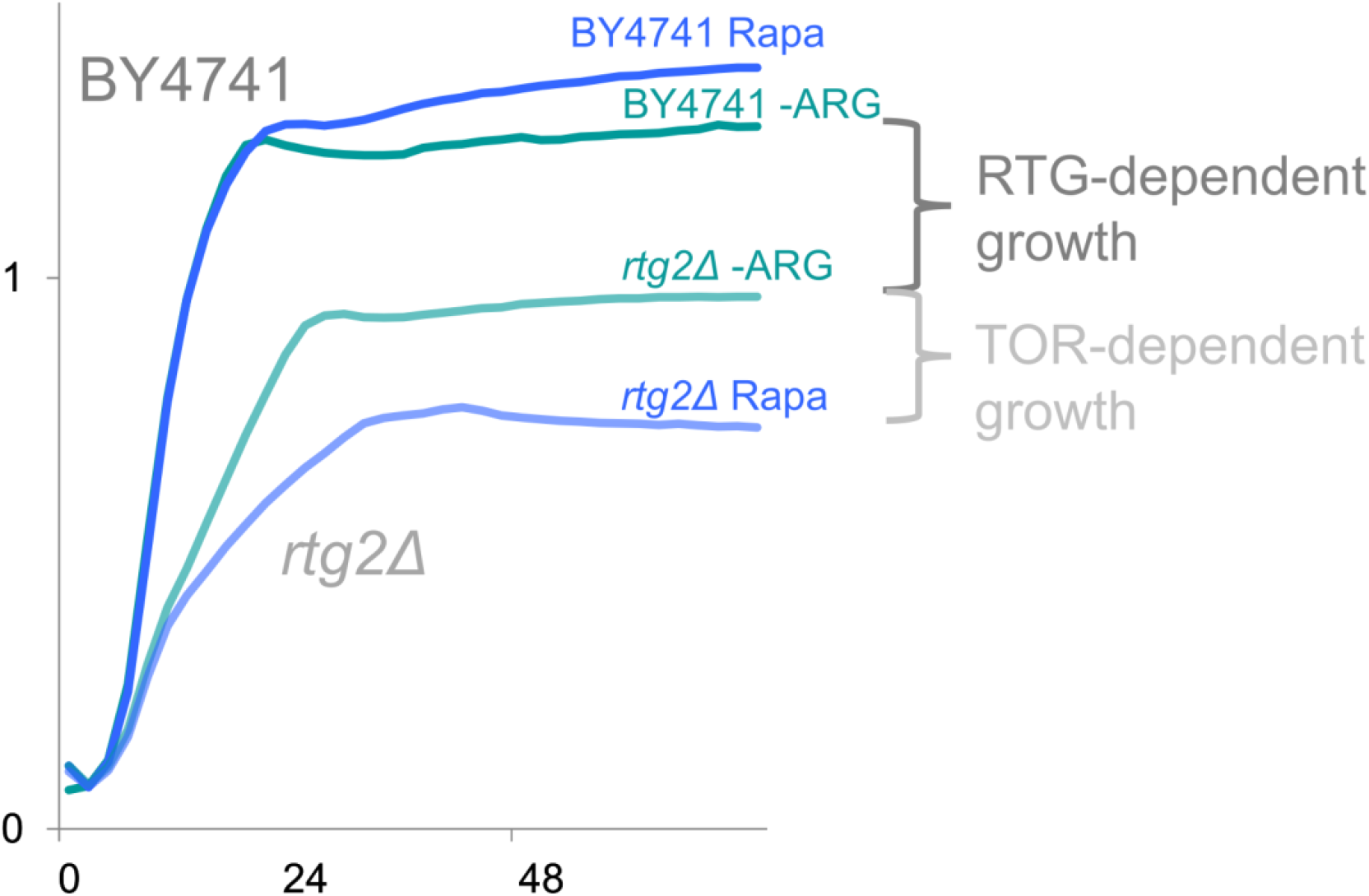
Both RTG activity and TOR activity are required for growth under arginine deprivation. Growth curves of BY4741 and *rtg2Δ* in liquid glucose (-)arginine (teal) and glucose (-)arginine + 0.025 μg/ml rapamycin (blue). X-axis = hours, Y-axis = OD_600_.

The work began with the observation that RTG single mutants exhibit growth deficiencies on glucose (-)arginine and are highly sensitive to canavanine - even more sensitive than petites. RTG mutants are also sensitive to thialysine, but don’t show as severe a growth inhibition on glucose (-)lysine. Doing the same experiment with a carbon source that induces a greater level of respiration restores growth on both (-)arginine and (-)lysine. Increasing respiration through this carbon source switch also improves tolerance to both canavanine and thialysine, although RTG mutants are still subject to the lactic acid effect on canavanine (Fig. 1). Taken together, this shows that the RTG pathway is required when glucose is present and yeast cells are subject to glucose repression. Relief of glucose repression by supplying an alternate carbon source, like galactose or lactic acid, rescues the growth deficiencies of RTG mutants grown in (-)arginine or (-)lysine, possibly through the transcriptional switch of *ACO1, CIT1, IDH1*, and *IDH2* to HAP control from RTG control, and increased TCA cycle runs that result from increased respiration. Increasing the number of TCA cycle runs would generate more αKG to fortify glutamate stores.

Our findings align with previous work showing that *rtg1Δ* growth is dependent on carbon source in minimal media - its growth improves as respiration is increased (11). It has also been previously reported that yeast grown in raffinose achieve resistance to acetic-acid-induced programmed cell death (AA-PCD) in an RTG-dependent manner; when grown on glucose, RTG mutants are highly sensitive to AA-PCD, but raffinose restores growth, although not to WT level (15). Raffinose is a carbon source that, like galactose, is partially fermented and partially respired. This complements our finding that under canavanine exposure, the need for RTG pathway activation is influenced by the fermentation/respiration ratio, as acetic acid is a byproduct of fermentation.

This prompted us to test if genetic relief of glucose repression via *MIG1* deletion could restore RTG mutant growth in a similar manner. On solid media, the carbon source effect equally rescues all single RTG mutants (Fig. 1), but *MIG1* deletion does not have an equal effect on the four tested RTG genes grown with glucose under arginine or lysine deprivation. *rtg1Δ* and *mks1Δ* are completely rescued by *mig1Δ* regarding both growth on glucose (-)arginine and growth with 1 μg/ml canavanine; *rtg2Δ* and *rtg3Δ* show a slight rescue in growth without arginine, but are still highly sensitive to canavanine on solid media (Fig. 2). Thus, although the four tested RTG mutants have a similar response to the carbon source switch, the different phenotypes of *RTG/MIG1* double mutants may indicate that the presence of glucose itself has a differential effect on the four RTG proteins of this study (Rtg1, Rtg2, Rtg3, and Mks1). One explanation is that certain genes are repressed by glucose, even when Mig1 is absent; such genes could possibly compensate for the loss of Rtg2/Rtg3. We currently aim to identify these genes using transcriptomics.

The central dogma of the RTG pathway is that Rtg2-dependent translocation of Rtg1/Rtg3 from the cytoplasm to the nucleus induces expression of RTG-target genes. Thus, in the *rtg2Δ* background, Rtg1/Rtg3 is not nuclear as it remains in the cytoplasm. Of course, this heterodimer is completely absent in *rtg1Δrtg3Δ*. Comparison of the growth curves of *rtg2Δ* and *rtg1Δrtg3Δ* in glucose (-)arginine and glucose (-)arginine + 1 μg/ml canavanine show that the two genetic backgrounds are not phenotypically equal under canavanine exposure. Yeasts that lack the Rtg1/Rtg3 heterodimer show less growth inhibition in liquid media than the *rtg2Δ* single mutant (Fig. 3C), suggesting that Rtg2 itself has an additional role in canavanine tolerance since RTG-target gene expression should be turned off in both strains. The need for Rtg2 is especially strong under glucose-repressing conditions, but it is still required when glucose repression is relieved through *MIG1* deletion. This is evident in the growth curves of the *RTG/MIG1* double mutants (Fig. 4). In liquid glucose (-)arginine + 1 μg/ml canavanine, *MIG1-* deletion gives significant rescue to not just *rtg1Δ* and *mks1Δ*, but also *rtg3Δ*, yet offers no growth improvement in the *rtg2Δ* background under the same conditions (Fig. 4B, Supp. Fig. 2). The RTG pathway is clearly required in the presence of glucose when even one amino acid (arginine or lysine) is lacking (Fig. 1), although *MIG1* deletion fully rescues growth on (-)lysine, but not on (-)arginine (Fig. 2). Taken together with the effect of carbon source on *rtg2Δ* growth, these results raise the possibility that Rtg2 is inactive under strictly respiratory conditions and the absence of a fermentable carbon source, and Rtg2 is required when a fermentable carbon source is present even if glucose repression is somewhat relieved. This explains why *rtg2Δ* has WT-level growth on lactic acid; under strict respiratory conditions, Rtg2 is not active in WT, which turns off the RTG pathway and HAP takes over (11). This hypothesis can be further tested by measuring Rtg2 expression or Rtg2/Mks1 binding in a fermentable and a non-fermentable carbon source (30).

Starting this work, our hypothesis was that RTG genes are required for the tolerance of toxic amino acid analogs (specifically those that are derived from glutamate) because the RTG pathway ensures sufficient glutamate is available for amino acid biosynthesis. Two of our findings complicate this idea: 1) *mks1Δ*, which should have the RTG pathway constitutively on, is also sensitive to canavanine and grows poorly on glucose (-)arginine, and 2) RTG mutants grown on glucose (-)lysine do not show equal growth defects from lysine deprivation as they do under arginine deprivation even though lysine biosynthesis also consumes glutamate and αKG. Each observation is discussed in detail below.

### Why do mks1Δ and rtg2Δmks1Δ show different phenotypes under arginine deprivation and canavanine exposure, and likewise under lysine deprivation and thialysine exposure?

In *mks1Δ* and *rtg2Δmks1Δ*, Rtg1/Rtg3 is constitutively nuclear and the RTG pathway is on, evident by high expression levels of RTG-target genes relative to WT and their glutamate prototrophy. A somewhat confusing observation is that on solid media, *mks1Δ* grows as poorly as the other RTG mutants. Yet, like the other RTG mutants, *mks1Δ’*s arginine deficiency is only present on glucose (Fig. 1). However, *rtg2Δmks1Δ* does show the expected phenotype on solid media. The observation that *rtg2Δmks1Δ* grows fine on solid media when arginine is missing supports prior work showing it is a glutamate prototroph; yet we don’t see WT-level growth on glucose (-)arginine (Fig. 1), and it has a growth curve similar to the petite strain (not WT) under canavanine exposure in liquid media (Supp. Fig. 2, Supp. Fig. 3), suggesting another factor involved in growth under these conditions has not been accounted for but is disproportionately affected in this background. Possibilities include mistargeting of arginine biosynthesis enzymes, canavanine-induced reduction of membrane potential, or altered TOR signaling. It’s important to note that Mks1 has pleiotropic roles in metabolism, including nitrogen catabolism (31), negative regulation of the cAMP-RAS pathway (32), and negative regulation of lysine biosynthesis, as it was first identified as *LYS80* (33).

Since TOR negatively regulates the RTG pathway through Mks1, we used rapamycin to pharmacologically inhibit TOR. In our experiments, both *mks1Δ* and *rtg2Δmks1Δ* show slight growth inhibition in response to rapamycin, even though it’s well-documented that RTG-target gene expression in *mks1Δ* and *rtg2Δmks1Δ* are unresponsive to rapamycin (3). Although *rtg2Δmks1Δ* shows a similar growth curve to *pif1Δ* under canavanine exposure, *rtg2Δmks1Δ* experiences growth inhibition by rapamycin just like the single RTG mutants and unlike *pif1Δ*. Thus, the TOR-dependent growth in these backgrounds suggests a signal between TOR and components of the RTG pathway that are not mediated through Mks1. One possibility is Lst8, which is a component of TORC1 and a known negative regulator of the RTG pathway. Being an essential gene, it was not included in this study. However, hypomorphic alleles of *lst8* that show constitutive activation of RTG-target gene expression suggest that Lst8 affects the RTG pathway in two ways - one upstream and one downstream of Rtg2. The site upstream of Rtg2 is thought to involve activity or assembly of the SPS (Ssy1-Ptr3-Ssy5) amino acid-sensing system, which affects cells’ ability to sense external glutamate (34). It is unclear how Lst8 negatively regulates the RTG pathway downstream of Rtg2 (35).

It has been hypothesized that since both Mks1 and TOR kinases are negative regulators of the RTG pathway, the Mks1-TOR complex may directly phosphorylate and inactivate Rtg3 (30). Additionally, a high-throughput screen examining a kinase and phosphatase interaction network shows a physical interaction between Kog1, an essential gene and subunit of TORC1, and Rtg3 (27). Thus, TOR potentially interacts with the RTG pathway through three different proteins: Mks1, Lst8, and Kog1. Supporting this notion, recent work shows that Kog1 regulates carbon assimilation and amino acid levels (28). We hypothesize that Rtg3 has an additional role besides being a nuclear transcription factor. This yet-to-be-characterized role is executed through interaction with Kog1 if there is sufficient glutamate, since Rtg3 should be cytoplasmic under sufficient glutamate levels. In *mks1Δ*, the Rtg1/Rtg3 transcription factor is located within the nucleus; this would prevent the interaction between Rtg3 and Kog1. Could this explain the growth deficiency on glucose (-)arginine and sensitivity of *mks1Δ* to canavanine? Perhaps, if this level of control overrides the benefits of constitutive expression of RTG-target genes. It is interesting to note that overexpression of Rtg3 (in the form of a GAL-induced plasmid) is lethal (36), suggesting that Rtg3 levels must be tightly regulated.

*MIG1* deletion rescues *mks1Δ* and *rtg1Δ* from canavanine toxicity in both solid and liquid conditions. Similar to how growth on lactic acid forces yeasts to respire, loss of *MIG1* releases the inhibition of respiratory genes. Increasing respiration and TCA cycle activity via *MIG1* deletion can explain the rescue of *mks1Δ, rtg1Δ*, and *rtg3Δ* in liquid canavanine culture, yet *MIG1* deletion offers no growth improvement in the *rtg2Δ* background under the same conditions (Fig. 4B, Supp. Fig. 2). TOR inhibition via rapamycin exposure further highlights the separation of function between the various RTG genes - when TOR is inhibited under canavanine exposure, *mks1Δ* shows full rescue by *MIG1* deletion; *rtg1Δ* shows a substantial rescue by *MIG1* deletion, although there is slight growth inhibition compared to *mig1Δ* and *mig1Δmks1Δ* (Fig. 5B, Supp. Fig. 2). The fact that *mig1Δ* doesn’t rescue *rtg2Δ* or *rtg3Δ* under canavanine + rapamycin suggests that these two genes have functions that can compensate partial inhibition of TOR. Following our hypothesis that Rtg3-Kog1 interaction is required for canavanine tolerance, it is possible that the metabolic remodeling provided by *MIG1* deletion in the *rtg1Δ* and *mks1Δ* backgrounds affects Rtg3 localization and/or phosphorylation status and consequently affects its interaction with Kog1.

Although Rtg3 has a potential physical interaction with TORC1, our results suggest that Rtg2 has the most important role in canavanine tolerance, especially when TOR is inactive; in canavanine, *rtg2Δ* is not rescued by *mig1Δ*, and *rtg2Δ* depends on TOR activity whether *MIG1* is present or not. This is strengthened by the observation that although *rtg2Δmks1Δ’s* growth curve in canavanine is most similar to the petite strain *pif1Δ*, unlike *pif1Δ, rtg2Δmks1Δ* cannot survive canavanine when TOR is inhibited by rapamycin (Supp. Fig. 2). Thus, the activation of RTG-target genes isn’t enough to override the absence of Rtg2 when TOR is inhibited by rapamycin.

### Why do RTG mutants show severe growth inhibition under arginine deprivation but not lysine deprivation?

We see very different growth on glucose (-)arginine versus glucose (-)lysine; specifically, RTG mutants show severe growth inhibition on solid glucose lacking arginine, but not lacking lysine (although there is some reduction in growth compared to WT and the petite). This is intriguing considering arginine and lysine biosynthesis both require αKG and glutamate. What defining characteristics does arginine have that lysine lacks?

The simplest explanation is that there are more pathways available for lysine production, or less barriers to production, in comparison with arginine; for example, unlike lysine biosynthesis, arginine biosynthesis relies heavily on mitochondrial transport because several arginine biosynthesis enzymes are encoded in the nucleus but function in the mitochondria (37). Alternatively, cellular arginine demands may be higher than lysine demands, possibly because arginine is used in protein synthesis at a higher rate. Three proteinogenic amino acids are encoded by six codons: arginine, leucine, and serine. The number of codons encoding for an amino acid correlate to the frequency of that amino acid being used within proteins (38); therefore, these three amino acids may be under extra regulation and required at higher amounts in order to make sure there are sufficient stores for protein synthesis (39). By comparison, lysine is only encoded by two codons. Another possibility is that arginine is specifically required for TOR activation in yeast. In mammals, SLC38A9 has been identified as an arginine sensor that activates TOR when there is sufficient arginine (40–43); conversely, arginine deprivation is sensed through mammalian CASTOR1 (a protein that is not part of the TOR complex), which inhibits TORC1 through an interaction with GATOR2 (44, 45). The GATOR complex is a multiprotein complex that responds to amino acid levels (9). The presence of arginine is not only sensed specifically by SLC38A9, but arginine disrupts this CASTOR1-GATOR2 interaction. Given the multiple arginine sensing mechanisms of mammalian TOR, it must be important for cells to properly sense and regulate arginine levels; given the highly conserved nature of TOR among different species, it’s possible that yeasts also contain mechanisms for TOR to specifically sense arginine. To the best of our knowledge, lysine does not have a specific effect on TOR activity in either mammals or yeast.

Although there is no identified equivalent of CASTOR1 in yeasts, Seh1-associated complex (SEAC) has been identified as the yeast equivalent of the mammalian GATOR complex (9). Thus, there may be some level of TOR inhibition in glucose (-)arginine due to arginine deprivation. When TOR is inhibited, the RTG pathway should be on to replenish αKG, glutamate, and, presumably, arginine. RTG mutants are incapable of doing this, which can account for their decreased growth compared to WT in glucose (-)arginine (Fig. 3). The phenotypic differences observed on glucose (-)arginine and glucose (-)lysine may be explained by a lack of lysine regulation over TOR - that is, lysine deprivation doesn’t inhibit TOR - plus, arginine is supplemented in glucose (-)lysine media, so TOR would be activated, leading to downregulation of the RTG pathway. Thus, knocking out the RTG pathway wouldn’t have as large of an effect because the cell relies less on the RTG pathway in glucose (-)lysine. Again, this only appears to be the case under glucose repressing conditions. RTG mutants grow just fine on galactose (-)arginine and lactic acid (-)arginine (Fig. 1). This raises questions about the status of TOR when an alternative carbon source is available.

The central dogma of the RTG pathway states that RTG activation operates through Rtg2-dependent translocation of the heterodimer Rtg1/Rtg3 from the cytoplasm to the nucleus. To the best of our knowledge, this work shows for the first time that Rtg3 function can be separate from its role as a heterodimer with Rtg1. This work also strongly suggests a specific role for Rtg2 in canavanine tolerance. Our findings suggest that *RTG* genes may have additional regulatory functions that don’t operate through the classically understood RTG pathway; rather, under certain conditions (low-dose canavanine exposure under arginine deprivation), Rtg2 and Rtg3 require a functioning TOR when glucose is present - a concept that expands the understanding of the relationship between TOR and the RTG pathway.

Analysis of the downstream targets of the RTG pathway reveals that genes directly involved in αKG levels are required for growth on (-)arginine, (-)lysine, canavanine, and thialysine (Fig. 6). Additionally, *IDH/CIT* double mutants show extra sensitivity to thialysine. This is likely due to the fact that both *CIT2* and lysine biosynthesis support peroxisome function (46, 47). Growth patterns of RTG-target gene mutants support our initial hypothesis that maintaining αKG levels is important for arginine and lysine biosynthesis. However, our results show that under arginine deprivation, loss of a single RTG gene causes greater growth inhibition than loss of two RTG-target genes; conversely, loss of a single RTG gene causes less growth inhibition under lysine deprivation, and instead loss of two RTG-target genes has a greater inhibitory effect on growth if one of those genes is *IDH1* or *IDH2* (Compare Fig. 1 to Fig. 6). What remains to be explored is what effect, if any, elevated D2HG levels have under toxic amino acid analog exposure. D2HG is an oncometabolite that can be mistakenly used by αKG-requiring enzymes, many of which participate in DNA maintenance. A hallmark of certain highly aggressive cancers is a mutated *IDH1/2*, which encodes for cytosolic human isocitrate dehydrogenase (48–50). Instead of converting isocitrate to αKG, these gain-of-function mutants convert αKG to D2HG – the reverse reaction of yeast Dld3 (51–53). In fact, arginine deprivation therapy itself, along with canavanine exposure, has shown potential as an anticancer therapy, especially against solid tumors (54–56). The work encompassed here may offer insight into the optimization of such therapies.

