## Supplementary figures and images for "Checks and balances of the RTG pathway under arginine deprivation and canavanine exposure in *Saccharomyces cerevisiae*"

### Supplementary Table 1

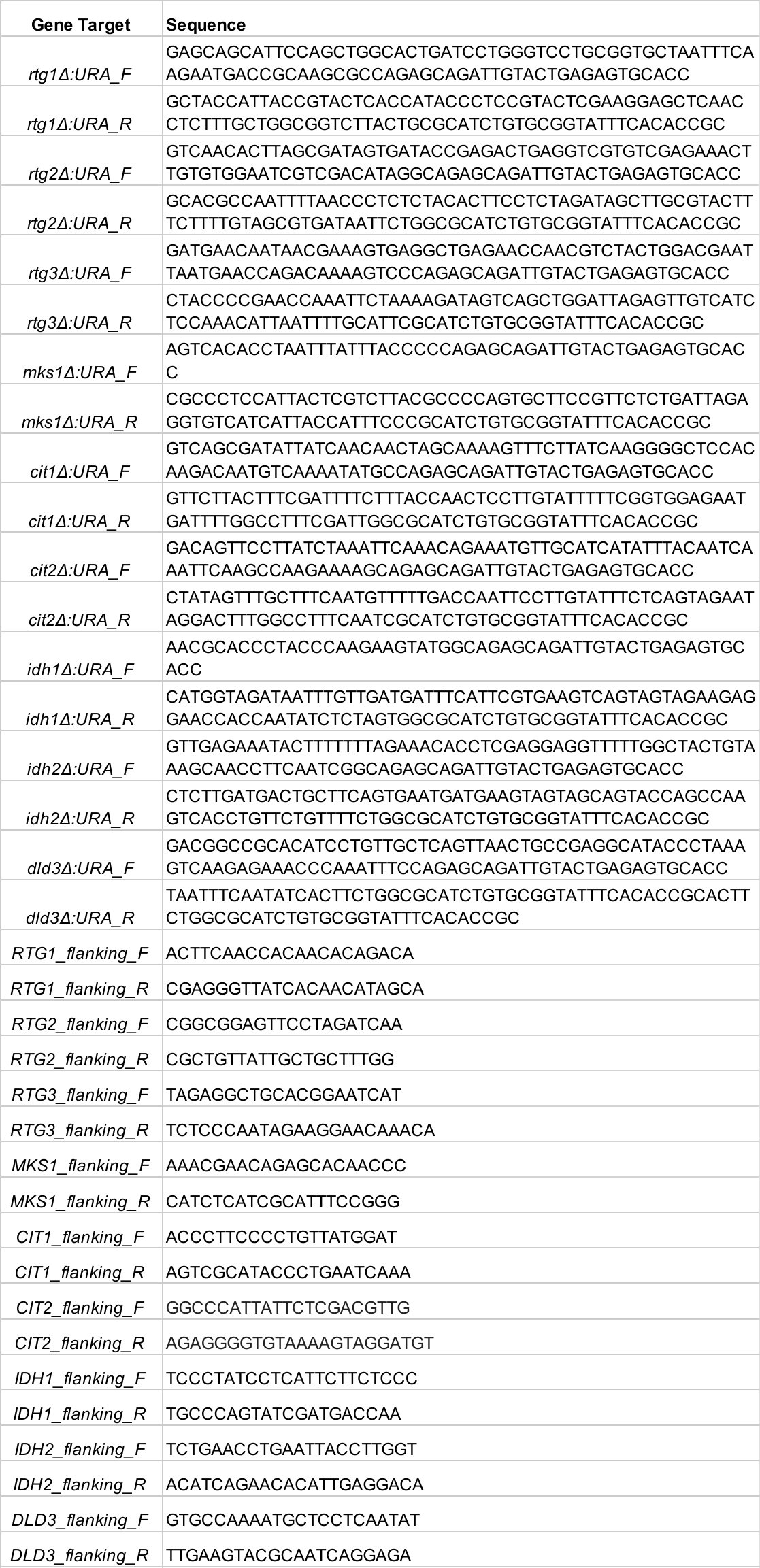
