## Supplementary Figures for "Checks and balances of the RTG pathway under arginine deprivation and canavanine exposure in *Saccharomyces cerevisiae*"

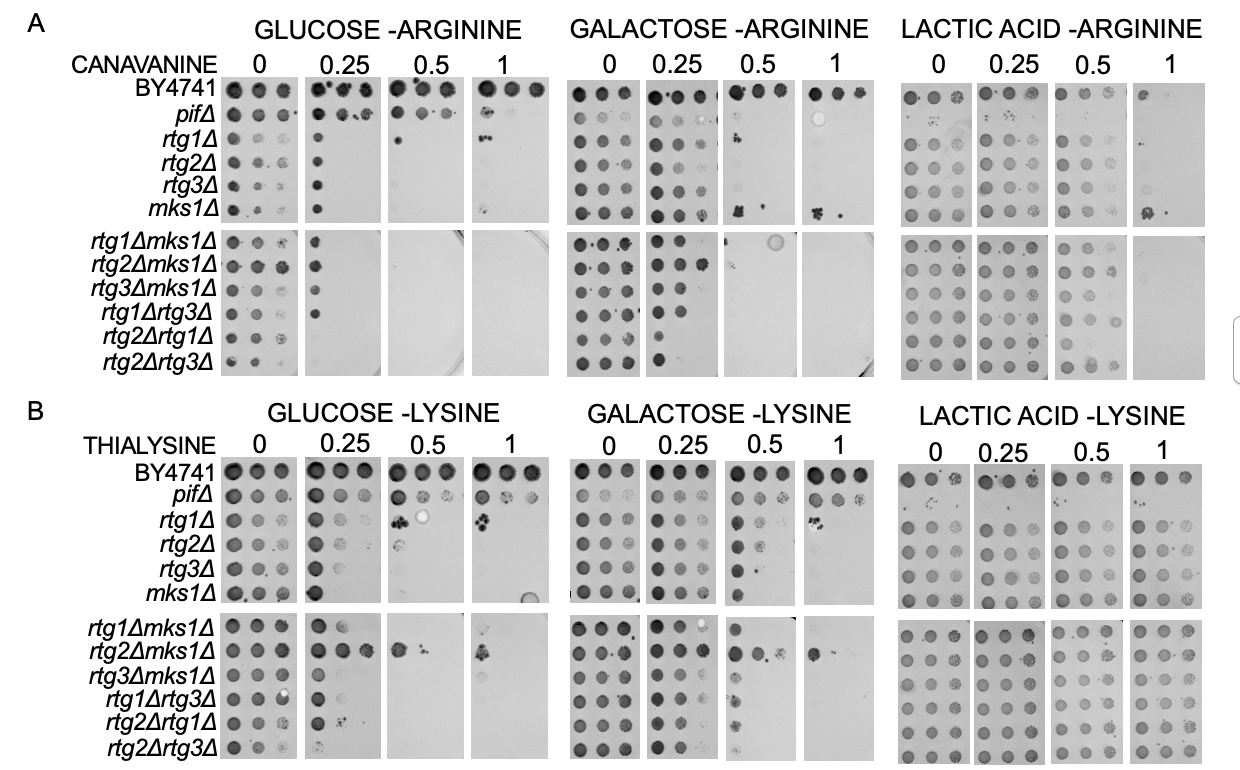

**Supplementary Figure 1. Alternative carbon sources improve growth of RTG mutants on (-)arginine and (-)lysine.** WT BY4741, petite strain *pif1Δ*, and *rtgΔ* single and double mutants grown on A) glucose (-)arginine, galactose (-)arginine, and lactic acid (-)arginine with the indicated amount of canavanine (µg/ml); and B) glucose (-)lysine, galactose (-)lysine, and lactic acid (-)lysine with the indicated amount of thialysine (µg/ml). A single colony was put in 100 ul of DDW, serially diluted, pronged onto the indicated media using a 6X8 metal pronger, and grown in 30^o^C. Growth after one week.

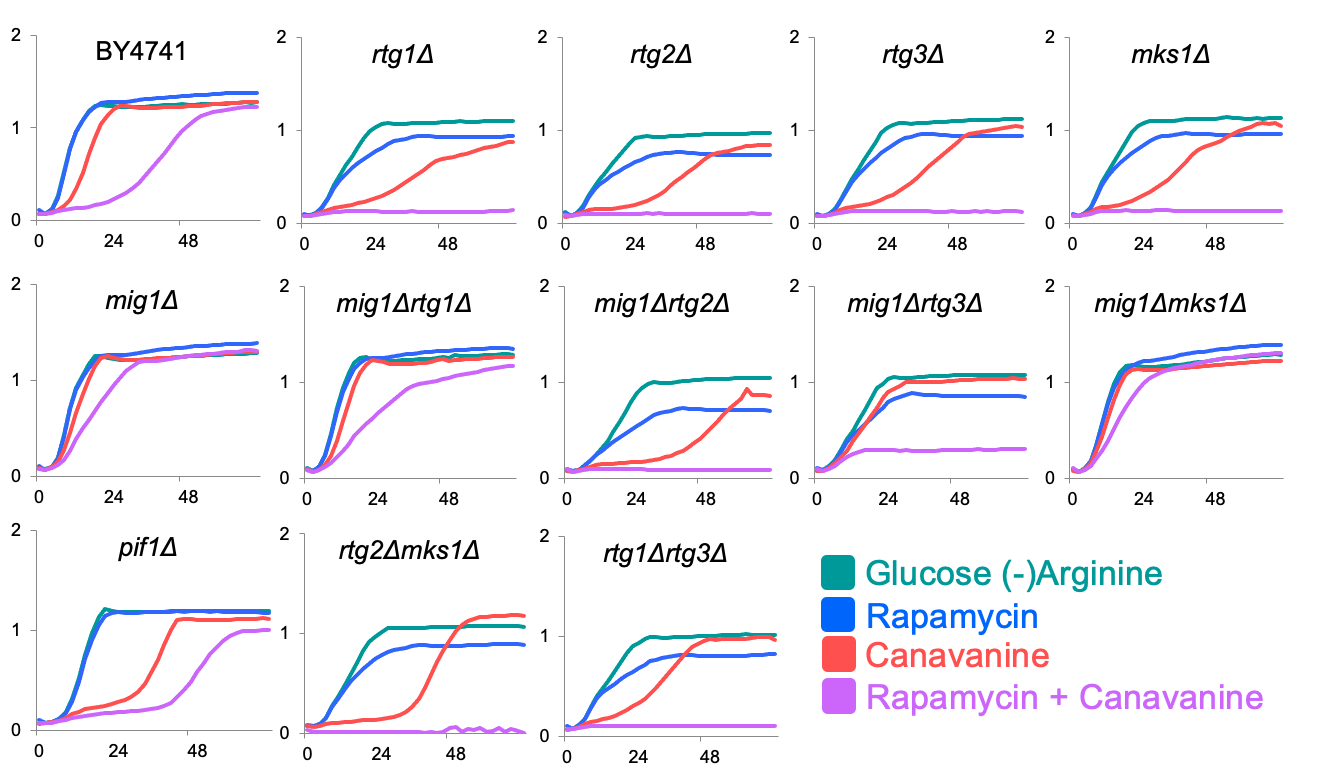

**Supplementary Figure 2. Growth dynamics of WT, *pif1Δ*, *mig1Δ*, and RTG mutants in glucose under arginine deprivation with and without rapamycin exposure and/or canavanine expoure.** Batch experiment showing growth of WT BY4741, *mig1Δ, pif1Δ*, *rtgΔ* single mutants, *mig1ΔrtgΔ* double mutants, *rtg2Δmks1Δ and rtg1Δrtg3Δ*. Growth in glucose (-)arginine (teal); glucose (-)arginine + rapamycin 0.025 µg/ml (blue); glucose (-)arginine + canavanine 1 µg/ml (red); and glucose (-)arginine + rapamycin 0.025 µg/ml + canavanine 1 µg/ml (purple). X-axis = hours, Y-axis = OD600.

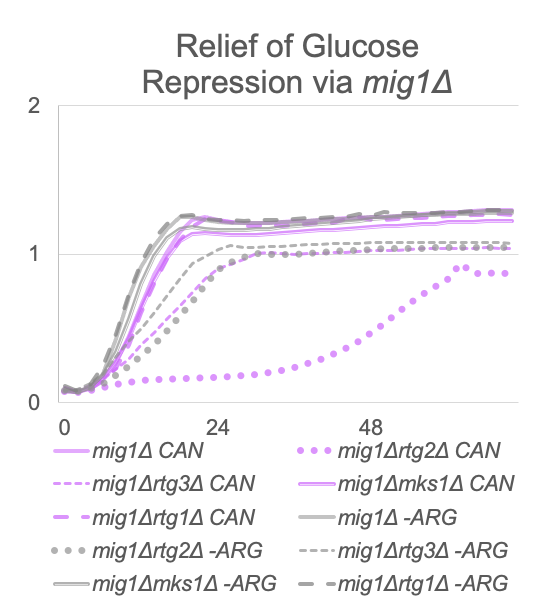

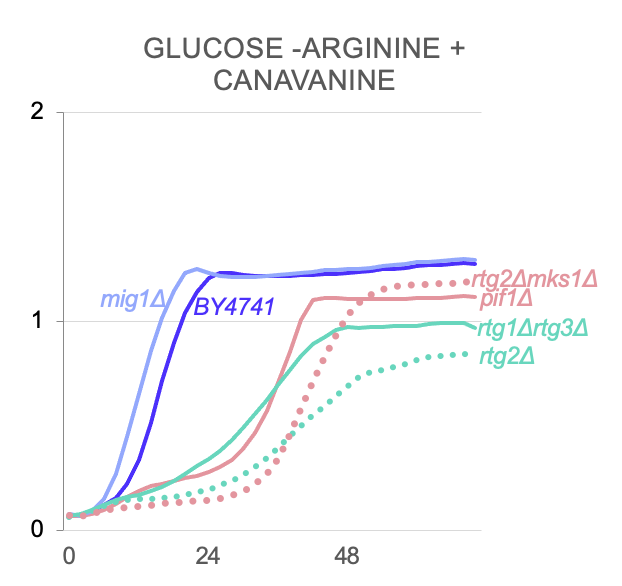

**Supplementary Figure 3. RTG-dependent growth and phenotypic differences in RTG mutants under canavanine exposure.** A) Growth curves of WT BY4741, *pif1Δ*, *mig1Δ, rtg2Δ, rtg1Δrtg3Δ,* and *rtg2Δmks1Δ* single mutants grown in liquid glucose (-)arginine + 1 µg/ml canavanine. RTG mutants and the petite strains *pif1Δ* spend extra time in lag phase relative to WT and *mig1Δ*. X-axis = hours, Y-axis = OD600.

**Supplementary Figure 4. *MIG1* deletion rescues RTG mutants except for *rtg2Δ* under canavanine exposure.** A) Growth curves of *mig1Δ* and *mig1ΔrtgΔ* double mutants grown in liquid glucose (-)arginine (gray) and glucose (-)arginine + 1 µg/ml canavanine (pink). X-axis = hours, Y-axis = OD600.
